# Characterization of the NSE6 subunit of the *Physcomitrium patens* PpSMC5/6 complex

**DOI:** 10.1101/2022.07.28.501545

**Authors:** E. Lelkes, M. Holá, J. Jemelková, B. Štefanovie, P. Kolesár, R. Vágnerová, K.J. Angelis, J.J. Paleček

## Abstract

Structural Maintenance of Chromosome (SMC) complexes are molecular machines ensuring chromatin organization at higher levels. They play direct roles in cohesion, condensation, replication, transcription and DNA repair. Their cores are composed of long-armed SMC, kleisin, and kleisin-associated KITE or HAWK subunits. Additional factors, like NSE6 within SMC5/6, bind to SMC core complexes and regulate their activities. To characterize the NSE6 subunit of moss *Physcomitrium patens*, we analyzed its protein-protein interactions and *Ppnse6* mutant phenotypes. We identified a previously unrecognized sequence motif conserved from yeast to humans within the NSE6 CANIN domain that is required for interaction with its NSE5 partner. In addition, the CANIN domain and its preceding sequences bind and link SMC5 and SMC6 arms, suggesting its role in SMC5/6 dynamics. Both *Ppnse6dCas9_3* and *Ppnse6KO1_47* mutant lines exhibited reduced growth and developmental aberrations. These mutants were also sensitive to DNA-damaging drug bleomycin and lost a significant portion of rDNA copies, suggesting conserved architecture and functions of SMC5/6 complexes across species.

## Introduction

Structural Maintenance of Chromosome (SMC) complexes are molecular machines driving chromatin organization at higher levels (Davidson & Peters, 2021; Uhlmann, 2016). They play key roles in cohesion, condensation, replication, transcription and DNA repair. Three bacterial (SMC/ScpAB, MukBEF, MksBEF), two archeal (canonical SMC/ScpAB and SMC5/6-related), and three eukaryotic (cohesin, condensin and SMC5/6) SMC complexes have been described (Mascarenhas *et al*, 2002; Petrushenko *et al*, 2011; Uhlmann, 2016; Yoshinaga & Inagaki, 2021). Their cores are composed of long-armed SMC, kleisin and kleisin-associated (KITE or HAWK) subunits. Additional factors are required for loading, targeting or regulation of SMC core complexes at specific chromatin sites (Litwin & Wysocki, 2018).

The SMC subunits are primarily built of head ATPase domains (formed by combined amino- and carboxyl-termini), long anti-parallel coiled-coil arms, and hinges (situated in the middle of the peptide chain; (Burmann *et al*, 2017; Diebold-Durand *et al*, 2017; Gligoris & Löwe, 2016; Hassler *et al*, 2018; Nasmyth & Haering, 2005)). Two SMC molecules form stable dimers via hinge domains at one end, and without ATP their arms align to rod-like structures. Binding of ATP molecules links the ATPase head domains at the opposite end and promotes the formation of the ring-shaped complex. ATP binding-hydrolysis cycle drives ring-to-rod dynamic changes and promotes DNA translocation or loop extrusion (Davidson *et al*, 2019; Ganji *et al*, 2018; Gruber, 2018; Wang *et al*, 2018).

The kleisin subunits bind “neck” and “cap” parts of SMC dimers and keep their head domains interconnected. In prokaryotic SMC and eukaryotic SMC5/6 complexes, kleisins interact with KITE (Kleisin-Interacting Tandem winged-helix Element) subunits (Palecek & Gruber, 2015), while cohesin and condensin complexes contain HAWK (HEAT-Associated With Kleisin) subunits (Wells *et al*, 2017). The SMC head-kleisin-KITE (or HAWK) form a compartment which anchors DNA during loop extrusion (Shaltiel *et al*, 2022; Yu *et al*, 2022). Uniquely, in addition to core SMC-kleisin-KITE subunits (called SMC5/SMC6-NSE4-NSE1/NSE3), SMC5/6 complexes contain highly conserved essential NSE2 SUMO-ligases (Andrews *et al*, 2005; Zhao & Blobel, 2005) which bind to SMC5 arms (Duan *et al*, 2009).

Additional NSE5 and NSE6 regulatory subunits interact with both SMC5 and SMC6 arms. Although their amino acid sequences are poorly conserved across the species, their primary function is to target SMC5/6 to the DNA damage sites (Oravcova *et al*, 2019; Oravcová & Boddy, 2019; Räschle *et al*, 2015). In the human HsNSE6, we recently identified a new CANIN (for Coiled-coil SMC6 And NSE5 INteracting) domain (Adamus *et al*, 2020). To further explore the conservation of this domain, we tracked down its sequence homology to lower plants and characterized its most conserved motif required for binding to its NSE5 partner. Using the lower plant *Physcomitrium patens* moss model organism, we demonstrated the conserved nature of the NSE6 subunit within the SMC5/6 complex.

## RESULTS

### *Physcomitrium patens* PpNSE6 CANIN domain interacts with PpNSE5

Recently, we characterized the human HsNSE6/SLF2 CANIN domain by showing its binding to the other HsSMC5/6 subunits (Adamus *et al*., 2020). To explore the conservation of these CANIN domain interactions, we tracked down the CANIN domain sequence homology to lower plants (PFAM ID number PF14816; (Mistry *et al*, 2021)) and selected model organism moss *Physcomitrium patens* PpNSE6 for further protein-protein interaction analysis. First, we prepared both N- and C-terminally truncated versions of Gal4AD-PpNSE6 and tested them against full-length (FL) Gal4BD-PpNSE5(1-526) construct in the yeast two-hybrid (Y2H) assay (Fig. 1). Deletion of the N-terminal 75 and 130 amino acids of PpNSE6 did not disturb its binding to PpNSE5. However, further deletion of the N-terminal 180 amino acids completely abolished PpNSE6 binding to PpNSE5 (Fig. 1B, lanes 1-4). Similarly, deletion of the C-terminal 110 amino acids of PpNSE6 did not abrogate its binding to PpNSE5, while further cutting into the CANIN domain (deletion of the last 150 amino acids) abolished it completely (Fig. 1B, lanes 5 and 6), suggesting the essential and conserved role of the CANIN domain (aa130 – 370) for PpNSE5-PpNSE6 interaction.

**Figure 1:**
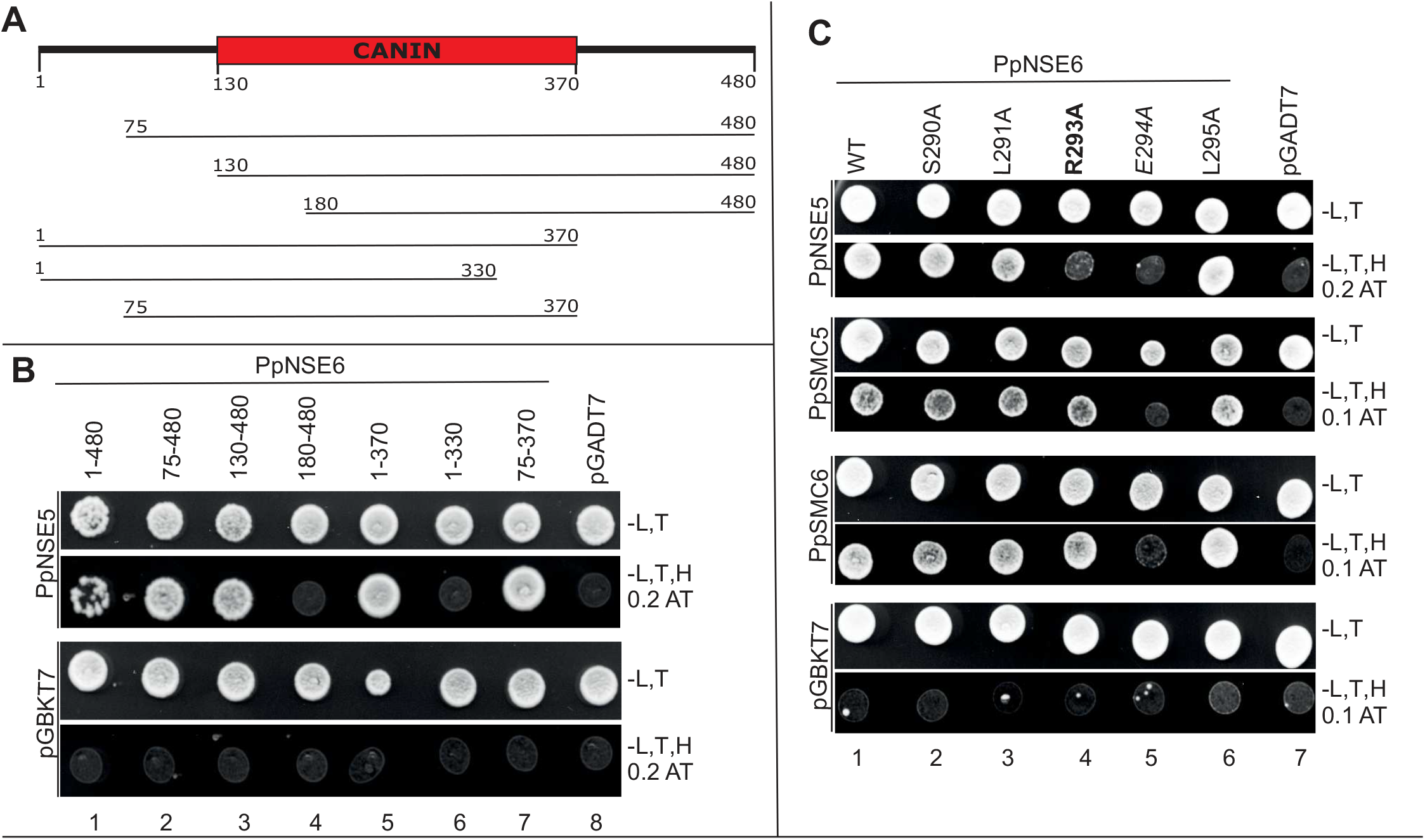
Detail analysis of the NSE5-NSE6 interaction. **(A)** Schematic representation of the different PpNSE6 constructs. The CANIN domain (aa130 – 370) is shown in red. **(B)** The indicated Gal4AD-PpNSE6 constructs were tested in Y2H for binding to the Gal4BD-PpNSE5(aa1-526). The intact CANIN domain was essential for the PpNSE5-PpNSE6 interaction. **(C)** Gal4AD-PpNSE6(1-370) mutants were tested in Y2H for binding to Gal4BD-PpNSE5(aa109-526), Gal4BD-PpSMC5(aa280-790), and Gal4BD-PpSMC6(aa226-955) constructs. The Ppnse6/R293A mutation (bold) specifically disturbed the PpNSE6-PpNSE5 interaction, while the Ppnse6/E294A mutation (italic) affected protein structure. All Y2H protein-protein interactions were scored by the growth of the yeast PJ69 transformants on the plates without Leu, Trp, His (-L,T,H), and with indicated concentration of 3-Amino-1,2,4-triazole (AT). Control plates were lacking only Leu and Trp (-L,T). Empty pGBKT7 and pGADT7 vectors were used as negative controls.

Using PSIPRED secondary structure predictions (McGuffin *et al*, 2000) and manually curated sequence alignments, we found previously unrecognized sequence homology between the CANIN domain of higher eukaryotes and budding yeast ScNSE6 sequence ((Pebernard *et al*, 2006; Zhao & Blobel, 2005); Suppl. Fig. S1A). In addition, modelling of the moss PpNSE6 structure (Jumper *et al*, 2021) provided a model similar to the yeast ScNSE6 structure (Taschner *et al*, 2021; Yu *et al*, 2021) over the most conserved part of the CANIN domain (Suppl. Fig. S1B). To explore the function of the most conserved motif, we mutated its amino acids to alanine (aa 290 to 295; Suppl. Fig. S1A) and tested them for binding to PpNSE5, PpSMC5 and PpSMC6 partners (Fig. 1C and see below). Of the five PpNSE6 mutations, R293A and E294A completely abolished the interaction with PpNSE5. While the R293A mutation disturbed only the PpNSE6-PpNSE5 interaction (Fig. 1C, lane 4), the ppnse6/E294A mutation affected the PpNSE6-PpSMC5 and PpNSE6-PpSMC6 interactions as well (lane 5). As ppnse6/E294A protein levels are normal (Suppl. Fig. S1D), we suggest that this mutation most likely affects the PpNSE6 structure. In conclusion, the residue Ppnse6/R293 specifically mediates the PpNSE6-PpNSE5 interaction and corresponds to the yeast ScNSE6/R318, which makes contact with ScNSE5 in crystal (PDB: 7OGG; (Taschner *et al*., 2021)) and cryoEM structures (PDB: 7LTO; (Yu *et al*., 2021)). Altogether, our data suggest a conserved interaction mode of NSE5-NSE6 across species.

### N-terminal and CANIN regions of PpNSE6 are necessary for binding to PpSMC5 and PpSMC6

The human HsNSE6/SLF2 CANIN domain was essential not only for binding to HsNSE5/SLF1, but also for the interaction with HsSMC6 (Adamus *et al*., 2020). Accordingly, the CANIN-containing Gal4AD-PpNSE6(aa1 – 370) Y2H construct bound both Gal4BD-PpSMC5 and Gal4BD-PpSMC6, while the CANIN-truncated PpNSE6(aa1 – 330) fragment did not bind to any of them (Fig. 2A, lanes 1 and 2), suggesting that the CANIN domain indeed plays an essential role in the binding to the core SMC5 and SMC6 subunits. Interestingly, deletion of the first 75 amino acids preceding the CANIN domain disturbed these interactions, while the binding to PpNSE5 was preserved (compare Fig. 1B, lane 7 with Fig. 2A, lane 3). This data suggests that both CANIN and N-terminal regions of PpNSE6 are essential for the binding to both PpSMC5 and PpSMC6.

**Figure 2:**
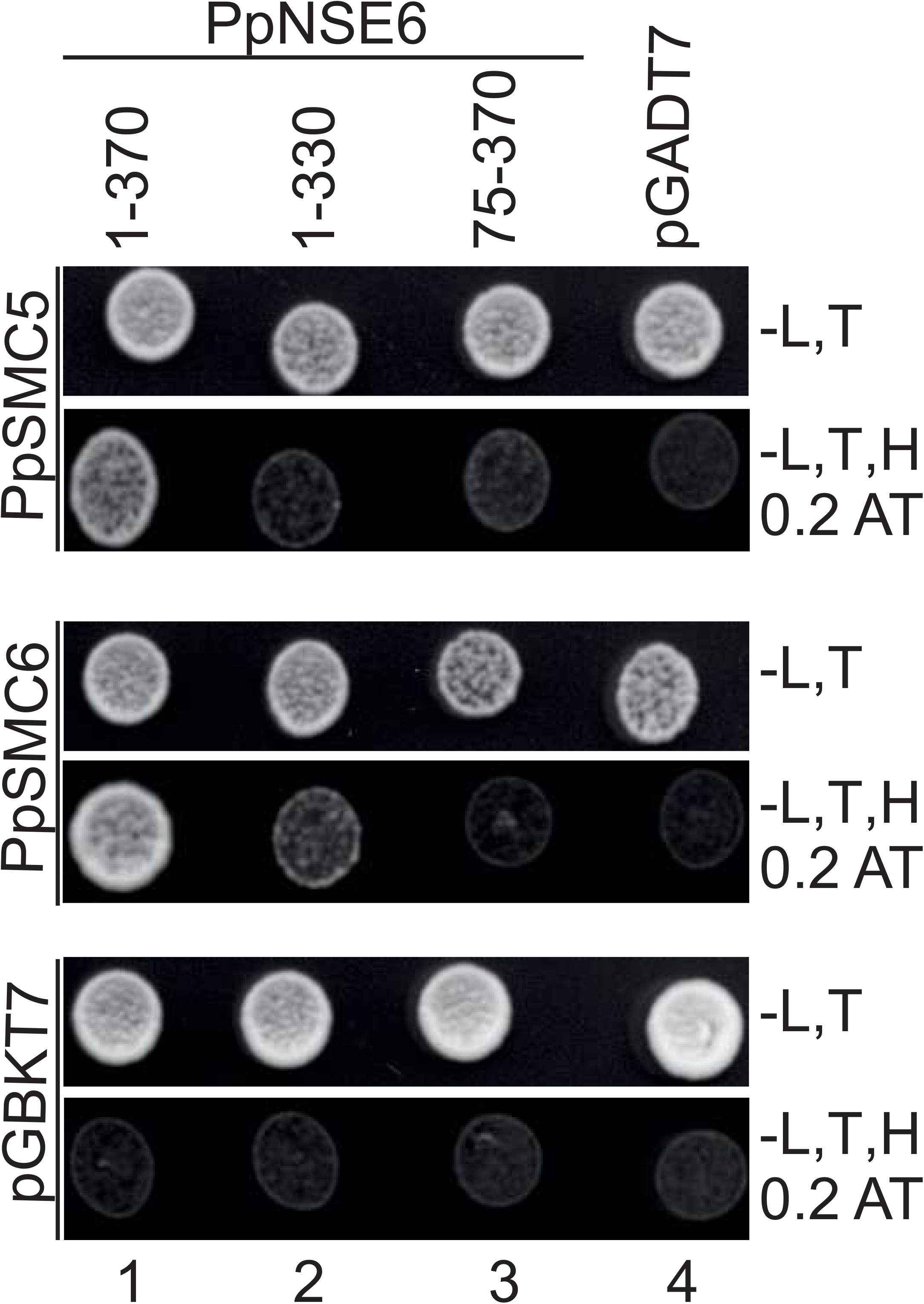
Analysis of PpNSE6 binding to PpSMC5 and PpSMC6. The indicated Gal4AD-PpNSE6 constructs were tested in Y2H for protein-protein interactions with Gal4BD-PpSMC5(aa201 - 890) (top panel) and Gal4BD-PpSMC6(aa226 - 955) (middle panel). The protein-protein interactions were scored in the same way as in Fig. 1.

### PpNSE6 binds PpSMC5 and PpSMC6 arms

Recently, we and others showed that the Nse6 subunits might link SMC arms (Adamus *et al*., 2020; Gutierrez-Escribano *et al*, 2020; Palecek *et al*, 2006; Taschner *et al*., 2021; Yu *et al*., 2021). Therefore, we prepared different fragments of both PpSMC5 and PpSMC6, containing hinge and coiled-coil segments of different lengths (Fig. 3). The Gal4BD-PpSMC5(aa201 – 890) and Gal4BD-PpSMC5(aa280 – 790) constructs bound Gal4AD-PpNSE6(aa1 – 370), while the shortest Gal4BD-PpSMC5(aa360 - 710) fragment did not (Fig. 3A, top panel, lanes 2 – 4). In the control experiment, these hinge-containing PpSMC5 constructs were able to bind to the PpSMC6 hinge-containing fragment (Fig. 3A, middle panel, lanes 2 – 4) via hinge-hinge interactions. These results suggest the essential role of PpSMC5 coiled-coil arm sequences aa280-360 and aa710-790 for the binding to PpNSE6.

**Figure 3.**
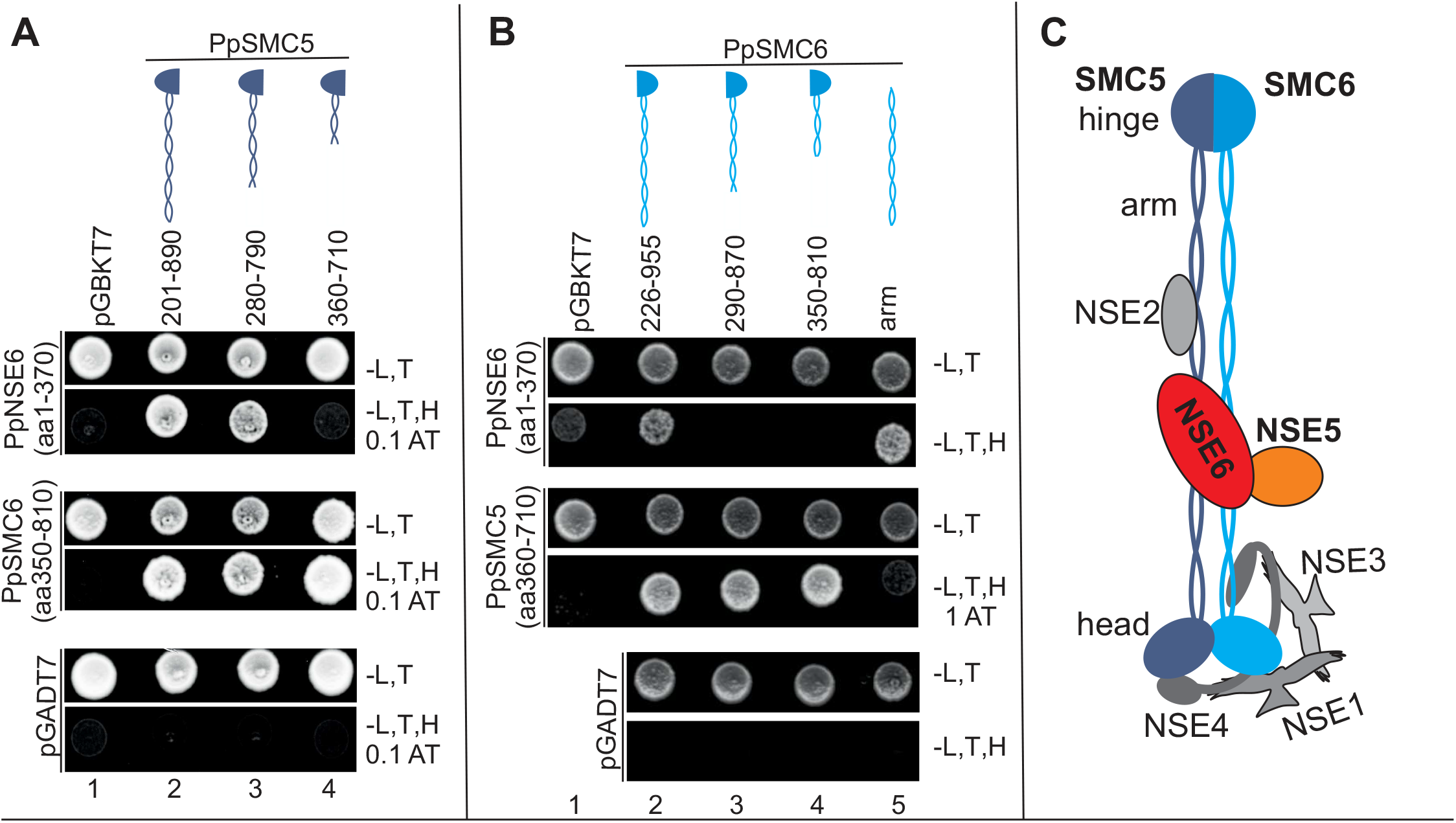
PpNSE6 binds to PpSMC5 and PpSMC6 arms. The Gal4AD-PpNSE6(aa1 - 370) binding to the indicated Gal4BD-PpSMC5 **(A)** and Gal4BD-PpSMC6 **(B)** constructs (top panels) was analyzed by the Y2H system. The binding of the indicated SMC constructs to partner hinge-containing fragments (Gal4AD-PpSMC5(aa360-710) and Gal4AD-PpSMC6(aa350-810), respectively) was used as a positive control (middle panels). Further details as in Fig. 1. **(C)** Model of the moss PpSMC5/6 complex. The N- and C-terminal halves of PpSMC proteins fold back at the hinge to form a typical SMC structure with composite arm and head domains. The NSE6 subunit links both SMC arms.

Similarly, the longest Gal4BD-PpSMC6(aa226 – 955) construct was able to bind Gal4AD-PpNSE6(aa1 – 370) while the shorter Gal4BD-PpSMC6(aa290 - 870) and Gal4BD-PpSMC6(aa350 - 810) fragments were not (Fig. 3B, top panel, lanes 2 – 4). These Y2H fragments were functional as they could bind to the Gal4AD-PpSMC5(aa360 - 710) hinge-containing fragment (Fig. 3B, middle panel, lanes 2 – 4; (Alt *et al*, 2017; Sergeant *et al*, 2005)). These data suggest the essential role of PpSMC6 coiled-coil arm sequences aa226-290 and aa870-955 for the binding to PpNSE6. Therefore, we made an arm construct without the hinge domain (aa226-510 fused to aa654-955) and showed its binding to PpNSE6 (Fig. 3B, top panel, lane 5). These results suggest that the PpSMC6 arm is sufficient while neither its hinge nor its head domain is required for the PpSMC6-PpNSE6 interaction. Altogether, the moss PpNse6 subunit binds PpNSE5 and links both SMC arms (Fig. 3C), similar to other organisms.

### Generation and analysis of moss *Ppnse6* mutants

Most SMC5/6 subunits are essential for the survival in most eukaryotic organisms, including plants (Diaz & Pecinka, 2018). Therefore, we first prepared *P. patens* mutant lines with reduced transcript levels using inactive dCas9 (Holá *et al*, 2021) to ensure their viability. We targeted the first exon of *PpNSE6* with dCas9-sgRNA and obtained stably transformed lines. The sensitivity of *Ppnse6dCas9* lines was scored as the growth of spotted explants treated 1 hr with 10 μg/ml bleomycin. The most sensitive lines were further propagated and analyzed for *PpNSE6* transcript levels by quantitative reverse transcription PCR (qRT-PCR). The *Ppnse6dCas9_3* line with the lowest transcript level (46% of wild-type level), was used further in this study.

To find out whether the NSE6 subunit is essential for *P. patens* viability, we next generated *nse6KO* STOP codon knock-in mutant. In the *PpNSE6* open-reading frame, the STOP codon was inserted via CRISPR/Cas9 homology-directed repair into the site of amino acid 9. We identified two lines with correctly integrated STOP codons, *Ppnse6KO1_35* and *Ppnse6KO1_47*. Both lines exhibited very similar morphology and growth response to bleomycin treatment. Nevertheless, the qRT-PCR showed that mutations do not affect the *PpNSE6* transcript levels. For further experiments, we chose the *Ppnse6KO1_47* line with a slightly more pronounced phenotype.

Both mutant lines, *Ppnse6dCas9_3* and *Ppnse6KO1_47*, exhibited reduced growth on a standard BCDAT medium. The protonemal growth was assessed as a fresh weight of explants 3 weeks after planting on Petri dishes with the drug-free BCDAT medium (Goffová *et al*, 2019). The growth rates were reduced by 30% in *Ppnse6dCas9_3* and 54% in *Ppnse6KO1_47* compared to wild-type (WT; Fig. 4A). Besides the reduced growth, both lines also manifested developmental aberrations, particularly inhibited formation of gametophores (Fig. 4B). Microscopic analysis of *Ppnse6dCas9_3* line shows predominant chloronemal filament growth with a reduced transition to caulonemall cell type which is indispensable for protonema side branching and gametophore formation (Prigge & Bezanilla, 2010). Propidium iodide (PI) staining of 10-days-old protonema showed that the rarely occurring apical caulonemal cells are often defective, exhibiting premature senescence leading to cell death in *Ppnse6dCas9_3* (Fig. 4C). In 10-days-old *Ppnse6KO1_47* protonema, the formation of caulonemal cells was inhibited completely, and the chloronemata showed deformed cell shapes.

**Figure 4.**
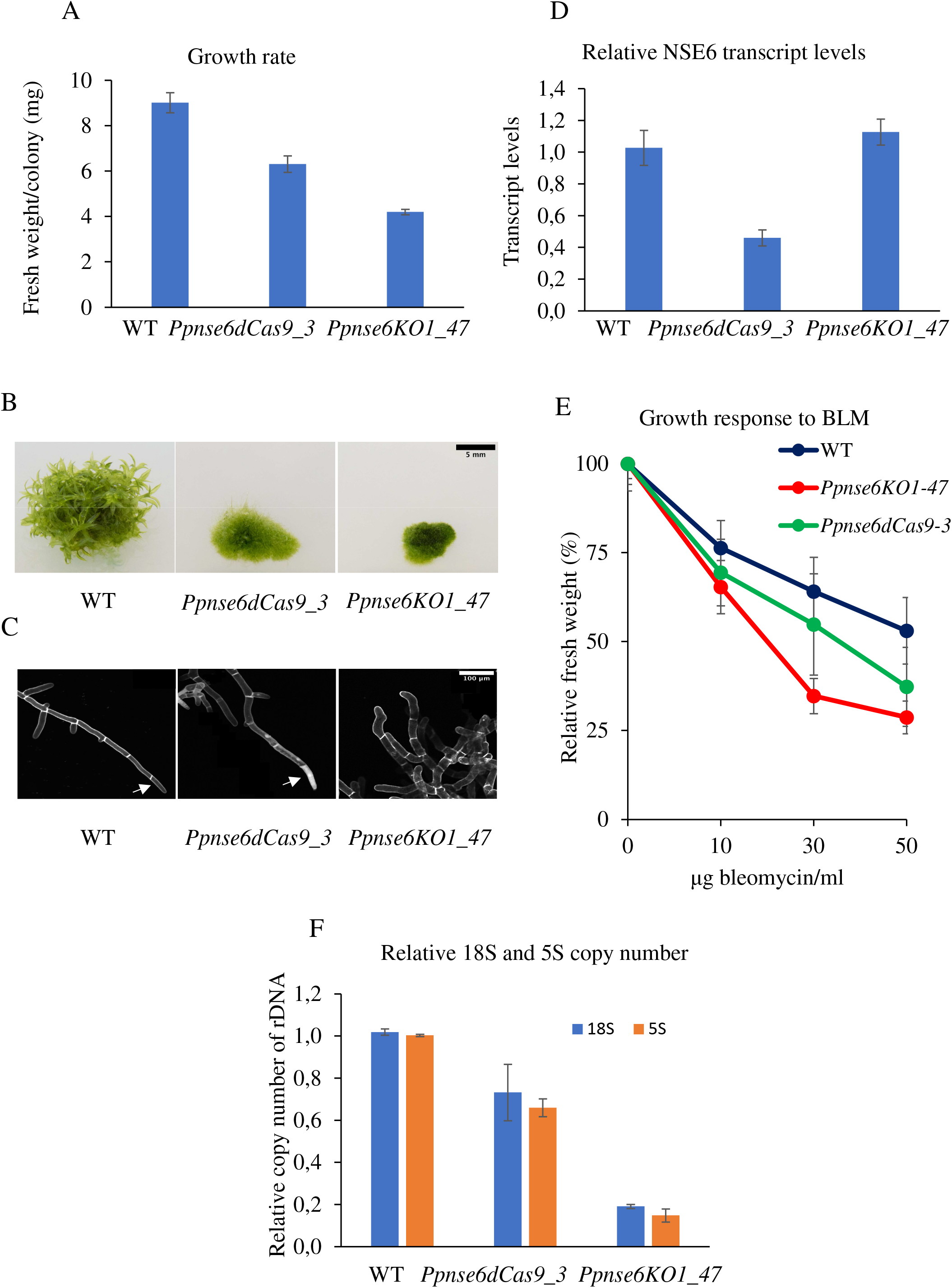
Characterization of moss *Ppnse6* mutant lines. (**A**) The growth rates of WT and mutant lines measured as fresh weight of 3-weeks-old untreated plants. (**B**) Morphology of 1-month-old colonies of WT and *Ppnse6* mutant lines grown on BCDAT medium (without bleomycin treatment). (**C**) 10-days-old protonema stained with propidium iodide. The apical caulonemal cells are indicated by arrows in WT and *Ppnse6dCas9_3*. In *Ppnse6KO1_47*, the transition to caulonemal cells is inhibited. (**D**) Relative levels of *PpNSE6* transcript and relative loss of 18S and 5S copies (**F**) in the *Ppnse6dCas9_3* and *Ppnse6KO1_47* mutant lines were measured by qPCR in three biological replicates and normalized to WT values. Error bars represent SD. (**E**) The growth response of WT and *Ppnse6* plants treated for 1 hr with 10, 30 and 50 µg/ml bleomycin. The explants were inoculated on a BCDAT medium and grown under standard conditions for three weeks. The mean weight of treated explants was normalized to the weight of untreated explants (set as a default 100%).

Growth-response to double-strand break (DSB) damage induced by radiomimetic drug bleomycin was used for characterization of defects in DNA repair processes of *Ppnse6dCas9_3* and *Ppnse6KO1_47* mutant lines (Holá *et al*., 2021). Consistently with the assumed role of SMC5/6 in DSB repair, *Ppnse6dCas9_3* and *Ppnse6KO1_47* were sensitive to bleomycin, with growth reduced to 55% and 35% of untreated controls, respectively (Fig. 4E) after 30 μg/ml bleomycin treatment.

Furthermore, given the role of SMC5/6 in rDNA stability maintenance (Peng *et al*, 2018; Torres-Rosell *et al*, 2005; Torres-Rosell *et al*, 2007), changes of rDNA copies were measured in genomic DNA of 7-days-old protonemata of *Ppnse6dCas9_3* and *Ppnse6KO1_47* by qPCR. The analysis revealed a statistically significant reduction of rDNA copies in both mutant lines. In *Ppnse6dCas9-3*, copy numbers of 18S rDNA were reduced to 73% and 5S rDNA to 66% (Fig. 4F). The loss of rDNA copy numbers in line *Ppnse6KO1_47* was even more severe with 18S rDNA copies reduced to 19% and 5S rDNA to 15% compared to WT. Notably, the rDNA copy numbers and bleomycin sensitivities (Figs. 4E and F) correlated with the severity of growth phenotypes (Figs. 4A-C). Altogether, our results suggest the conserved interactions and functions of PpNSE6 within SMC5/6 complexes.

## Discussion

This study shows the conserved character of the NSE6 CANIN domain binding to its NSE5 partner. Moss PpNSE6 bound its PpNSE5 partner via the CANIN domain (Fig. 1), similar to human HsNSE6 (Adamus *et al*., 2020). Within the CANIN domain, we identified a conserved sequence motif which mediates NSE6-NSE5 interactions across species, as seen in structures of budding yeast ScNSE6 (Taschner *et al*., 2021; Yu *et al*., 2021) and demonstrated for moss PpNSE6. Although the sequence homology of the NSE6 CANIN domain is not as high as that of the core SMC5/6 subunits (Hudson *et al*, 2011; Palecek *et al*., 2006; Yoshinaga & Inagaki, 2021), it is sufficient to mediate its functional interaction with NSE5 in organisms from yeast to human.

The NSE6 CANIN domain is also required for the SMC6 arm interaction of yeast, plants and human subunits (Fig. 2; (Adamus *et al*., 2020; Gutierrez-Escribano *et al*., 2020; Taschner *et al*., 2021; Yu *et al*., 2021). However, other than the above-identified conserved motif and additional N-terminally located amino acids are required for binding to SMC6 in yeast (Taschner *et al*., 2021) and moss (Fig. 2). In addition to SMC6 binding, the NSE6 subunits also bind SMC5 and connect these core subunits at their arms (Adamus *et al*., 2020). We hypothesized that this bridging of SMC5-SMC6 arms might negatively impact their dynamics, e.g. by limiting the opening of the SMC5/6 ring upon ATP binding (Adamus *et al*., 2020; Palecek *et al*., 2006). This kind of ATPase activity inhibition was observed for budding yeast complexes (Hallett *et al*, 2021; Taschner *et al*., 2021). Our new results suggest that this NSE6 inhibitory effect might be conserved in plants and humans as their NSE6 subunits link SMC5-SMC6 arms.

The NSE6-mediated inhibition of SMC5/6 ATPase activity suggests a regulatory role of NSE6 rather than a structural role. Consistent with this notion, the association of the NSE6 subunit with the SMC5/6 core complex has been implicated specifically in the localization to the DNA repair sites (Oravcova *et al*., 2019; Oravcová & Boddy, 2019; Räschle *et al*., 2015). Accordingly, the *S. pombe* spNSE6 gene is required for damage repair, but it is not essential for fission yeast cell survival (Pebernard *et al*., 2006), while the SMC5/6 core subunits are essential (Lehmann *et al*, 1995). Interestingly, we were able to generate viable *Ppnse6KO* lines (Fig. 4), while our attempts to prepare *Ppsmc6KO* lines failed (Holá *et al*., 2021). Nevertheless, the *Ppnse6* mutant lines exhibited severe phenotypes with reduced cell viability and high sensitivity to DNA damage, similar to other moss *Ppsmc5/6* mutant lines (Holá *et al*., 2021) or other plant mutants (Diaz & Pecinka, 2018). The reduced viability might stem from compromised ability to recover from (endogenous) DNA damage or rDNA copy number deficiency (Fig. 4; (Torres-Rosell *et al*., 2005; Torres-Rosell *et al*., 2007)) or other SMC5/6-related defects (Aragón, 2018), which lead to cell death when accumulated.

## Materials and Methods

### Yeast two-hybrid constructs and assays

The full-length PpNSE6 and fragments containing aa 1-330, aa75-480 and aa180-480 were cloned into *Nde*I-*Bam*HI digested pGADT7 using NEBuilder (New England BioLabs, USA) after PCR amplification from the template DNA (CM189) with primers specified in Suppl. Table ST1. PpNSE6 (aa1-370) and (aa130-480) were first cloned into the pGBKT7 *Nco*I-*Bam*HI sites using NEBuilder (primers specified in Suppl. Table ST1), then cleaved out with *Nco*I-*Bam*HI and ligated into *Nco*I-*Bam*HI sites of pGADT7. The PpNSE6 construct with aa75-370 was first cloned into the pGBKT7 *Nde*I-*Bam*HI sites with NEBuilder, then cleaved out with *Nde*I-*Bam*HI and ligated into the *Nde*I-*Bam*HI sites of pGADT7.

The pGBKT7-PpNSE5 (aa1-526) and (aa109-526) constructs were created by amplifying the PpNSE5 gene from cDNA and cloning it into the pGBKT7 *Nde*I-*Bam*HI sites (CM186) using NEBuilder (primers specified in Suppl. Table ST1).

The pGBKT7-PpSMC5(aa201-890) was created by amplifying the PpSMC5 gene from cDNA and cloning it into the pGBKT7 *Nde*I-*Eco*RI sites (MH1). The other fragments (aa280-790 and aa360-710) were amplified from MH1 using primers specified in Suppl. Table ST1 and cloned into pGBKT7 *Nde*I*-Bam*HI sites. The pGBKT7-PpSMC5(aa360-710) was cleaved by *Nde*I*-Bam*HI, and the fragment was inserted into the pGADT7 vector.

The pGBKT7-PpSMC6(aa226-955) was made by amplifying the PpSMC6 gene from cDNA and cloning it into the pGBKT7 *Nde*I-*Eco*RI sites (MH3). The other pGBKT7-PpSMC6 constructs containing the arms and the hinge domains (aa290-870 and aa350-810) were prepared by PCR amplification of MH3 using primers specified in Suppl. Table ST1 and cloned into *Nde*I*-EcoR*I sites of pGBKT7 with NEBuilder. The pGBKT7-PpSMC6 (aa350-810) was cleaved by *Nde*I*-EcoR*I and ligated into *Nde*I*-EcoR*I sites of pGADT7. The pGBKT7-PpSMC6 head-arm (aa1-510 + aa655-1095) construct (PEB197) was created in two steps. First, the N-terminal head-arm part (aa1-510) was amplified by PCR from cDNA and cloned into pGBKT7 (*Nde*I site). Second, the C-terminal head-arm part (aa655-1095) was PCR amplified from cDNA and inserted into the N-terminal construct (*Bam*HI site) created in the first step. Finally, the pGBKT7-PpSMC6 arm (aa226-510 + aa655-955) construct was prepared from the PpSMC6 head-arm (aa1-510 + aa655-1095) construct (PEB197) by PCR amplification (using primers EB259 and EB260) and insertion into *Nde*I-*Eco*RI site of pGBKT7.

The QuikChange Lightning Site-Directed Mutagenesis Kit (Agilent Technologies, USA) was used to create mutations in the pGADT7-PpNSE6(aa1 – 370) construct. The sequences of primers used for mutagenesis are listed in the Suppl. Table ST1.

The classical Gal4-based Y2H system was used to analyze the SMC5/6 complex interactions as described previously (Hudson *et al*., 2011). Briefly, pGBKT7 and pGADT7 constructs were co-transformed into the *Saccharomyces cerevisiae* PJ69–4a strain and selected on SD -Leu, -Trp plates. Drop tests were carried out on SD -Leu, -Trp, -His (with 0.1; 0.2; 0.3; 0.5; 1; 2; 3; 4; 5; 10; 15; 20; 30 mM 3-aminotriazole) plates at 28°C. Each combination was co-transformed at least three times, and at least three independent drop tests were carried out.

### Analysis of protein structures

The AlphaFold (Jumper *et al*., 2021; Varadi *et al*, 2022) and SwissModel (Waterhouse *et al*, 2018) tools were used to generate *in silico* protein models. All structures were visualized by the Pymol software (Schrodinger Inc., USA).

### Plant material and cultivation

The WT *P. patens* (accession (Hedw.) B.S.G.; (Rensing *et al*, 2008)) was used for the generation of *Ppnse6* mutants. The moss lines were cultured as a ‘spot inocula’ on BCD agar medium supplemented with 1 mM CaCl_2_ and 5 mM ammonium tartrate (BCDAT medium), or as lawns of protonema filaments by subculture of homogenized tissue on BCDAT agar overlaid with cellophane in growth chambers with 18/6 h day/night cycle at 22/18 **°**C (Cove *et al*, 2009).

### Construction and analysis of *PpNSE6* mutant lines

The vector for attenuation of *PpNSE6* transcription was constructed as expression cassettes of dCas9 driven by the maize ubiquitin promoter, a Hyg^R^ selection cassette and a specific guide RNA (sgRNA). sgRNA for *NSE6* was synthesized as two complementary oligonucleotides (pKA1355, pKA1356; Suppl. Table ST2) and targeted into the first exon in position +7 to +25. Entire construct was flanked by sequences homologous to neutral locus pp108 (Schaefer & Zrÿd, 1997), to improve stable integration into the genome. 30μg of DNA construct cleaved by *Bsa*I were delivered into protoplasts by PEG-mediated transformation as described in (Liu & Vidali, 2011). After five days of regeneration, the transformed protoplasts were transferred to a Petri dish with the BCDAT medium supplemented with 30 mg/l Hygromycin. After three rounds of selection, the transformants were considered stable. Transformants of *Ppnse6dCas9* were treated with 10μg/ml bleomycin supplied as Bleomedac inj. (Medac, Hamburg, Germany) for 1 hr, and planted as ‘spot inocula’ on BCDAT medium. After three weeks, the transformants with the most reduced protonemal growth were selected and analyzed by qRT-PCR for *PpNSE6* transcription.

Total RNA was isolated from 7-days-old protonemata with RNeasy Plant Mini Kit (Qiagen), treated with DNaseI (DNA-*free*™ DNA Removal Kit, Thermo Fisher Scientific) and reverse transcribed using qPCRBIO cDNA Synthesis Kit (PCR Biosystems). Diluted cDNA reaction mixtures were used for qRT-PCR analysis using the qPCRBIO SyGreen Mix Lo-ROX (PCR Biosystems) in Stratagene-MX3005P. Analysis was performed for three biological replicas (independently cultivated tissue) and in two technical replicates with Clathrin adapter complex subunit CAP-50 (Pp3c27_2250V3.1) as a reference gene (Kamisugi *et al*, 2016). The relative transcription of *PpNSE6* was calculated by the ΔΔCt method (Pfaffl, 2004).

The STOP-codon knock-in was introduced by homology-directed repair (HDR) after Cas9-induction of DSB within the *NSE6* locus and 40 bp double-stranded DNA donor template with the desired mutation. The donor template was composed of two complementary oligonucleotides (pKA1400 and pKA1401; Suppl. Table ST2) homologous to the first exon of *NSE6* locus and contained substitution of +25G to T (generating STOP codon at the site of 9^th^ amino acid) and +11GC to CG generating *Bsm*BI cleavage site. Gateway destination vector with Cas9 expression cassette (pMK-Cas9-gate) and kanamycin resistance, and entry vector containing the PpU6 promoter and sgRNA(pENTR-PpU6sgRNA-L1L2) were kindly provided by prof. Bezanilla (Mallett *et al*, 2019). sgRNA was synthesized as two complementary oligonucleotides (pKA1402 and pKA1403; Suppl. Table ST2). Four nucleotides were added to the 5′ ends of the oligonucleotides such that, when annealed, they create sticky ends compatible with *Bsa*I□linearized pENTR-PpU6sgRNA-L1L2. Cas9/sgRNA expression vector was generated using the Gateway LR reaction to recombine the entry vector pENTR-PpU6sgRNA-L1L2 with sgRNA spacer and destination vector pMK-Cas9-gate. DNA construct was co-transformed with donor template (annealed pKA1400 + pKA1401) into protoplasts by PEG-mediated transformation. All sgRNAs were designed in the CRISPOR online software using *P. patens* (Phytozome V11) and *S. pyogenes* (5′ NGG 3′) as the genome and PAM parameters, respectively. The protospacers with the highest specificity score were chosen.

After five days of regeneration, the transformed protoplasts were transferred to the BCDAT medium supplemented with 50 mg/l G418. After one week of selection, the G418 resistant lines were propagated. Crude extracts from young tissues of these lines were used for PCR amplification of genomic DNA around editing sites (primers pKA1451, pKA1452). The PCR products were cleavaged by *Bsm*BI and examined by DNA electrophoresis. The lines whose PCR product was cleaved were sequenced to confirm the correct introduction of mutations.

### Growth rate and Sensitivity Assays

Sensitivity to DNA damage was measured after the treatment with freshly prepared solutions of bleomycin sulphate. Protonemata were dispersed in a liquid BCDAT medium containing bleomycin for 1 hr. After treatment and rinse, the recovered protonemata were inoculated as eight explants per quadrant of a Petri dish with drug-free BCDAT agar without cellophane overlay and left to grow. The treatment effect was assessed after three weeks by weighing explants. The fresh weight of the treated explants was normalized to the fresh weight of untreated explants of the same line and plotted as % of ‘Relative fresh weight’. The growth rate of mutant lines was determined by weighing of untreated explants of mutant lines and their comparison with untreated explants of WT. In every experiment, the treatment was carried out in duplicate and experiments were repeated eight or ten times and statistically analyzed by Student’s t-test.

### DNA isolation and rDNA copy numbers analysis

DNA was isolated from 7-day-old protonemal tissue according to (Dellaporta *et al*, 1983). The DNA quality was checked and concentration was determined by electrophoresis in a 1% (w/v) agarose gel stained with ethidium bromide and using Gene Ruler 1 kb DNA Ladder (Thermo Scientific, Waltham, MA, USA) as standard.

qPCR for rDNA copy number analysis was performed in technical triplicates for three biological replicates of samples to analyze 18S rDNA (primer combination Pp18S_A and Pp18S_B) and 5S rDNA (primer combination Pp_5S_F and Pp_5S_R) normalized to ubiquitin as a reference gene (primer combination ubqFw and ubqRev; (Goffová *et al*., 2019)) using qPCRBIO SyGreen Mix Lo-ROX (PCR Biosystems) in Stratagene-MX3005P. The data were statistically analyzed using two-tailed unpaired Student′s t-test.

### Microscopic analysis

10-days-old protonemata were stained with 10µg/ml propidium iodide (PI) (Sigma-Aldrich) in liquid BCDAT medium, mounted onto a glass slide and analyzed by a Spinning disc (SD) microscope Nikon (Eclipse Ti-E, inverted) with Yokogawa CSU-W1 SD unit (50mm), on a Nikon Ti-E platform, Laser box MLC400 (Agilent) Zyla cMOS camera (Andor), Nikon Plan-Apochromat L 10x/0.45 objective.

## Abbreviations

CANIN,: Coiled-coil SMC6 And NSE5 INteracting;
DSB,: Double-Strand Break;
HAWK,: HEAT-Associated With Kleisin;
KITE,: Kleisin-Interacting Tandem winged-helix Element;
NSE,: Non-SMC Element;
PI,: propidium iodide;
qRT-PCR,: quantitative reverse transcription polymerase chain reaction;
SMC,: Structural Maintenance of Chromosomes;
WT,: wild-type;
Y2H,: Yeast Two-Hybrid

## Acknowledgements

We thank Marek Adamus for his technical assistance and critical reading of the manuscript. Funding from the Czech Science Foundation (project GA20-05095S) is gratefully acknowledged.

