## Supplementary data for "Characterization of the NSE6 subunit of the *Physcomitrium patens* PpSMC5/6 complex"

### Materials and methods

#### Protein expression analysis

Yeast cells were grown to OD~1 and lysed by incubation in 0.1 M NaOH for 5 min and boiling in SDS Laemmli buffer (62.5 mM Tris–HCl, 2% SDS, 5% β-mercaptoethanol, 10% glycerol, 0.002% bromophenol blue; (Kushnirov, 2000)). Samples were separated by 12% SDS-PAGE and immunoblotted with anti-Gal4AD (Sigma Aldrich - G9293) antibody.

Kushnirov, V.V. (2000). Rapid and reliable protein extraction from yeast. Yeast *16*, 857-860.

### Figure legends

**Suppl. Figure S1: (A)** Alignment of the core part of the CANIN domain with helical (H) segments from ScNSE6 PDB: 7LTO structure (above) and AlphaFold PpNSE6 prediction (below). The most conserved central helical motif is highlighted in blue and red, respectively (compare to panel **B**). The conserved basic contact residues are marked by a blue arrowhead. The NSE6 orthologs are from *Saccharomyces cerevisiae* (*Sc*), *S. pastorianus* (*Sp*), *Zygosaccharomyces bailii* (*Zb*), *Physcomitrium patens* (*Pp*), *Arabidopsis thaliana* (*At*), *Brasica napus* (*Bn*), *Xenopus leavis* (*Xl*), *Galus galus* (*Gg*), *Ornithorhynchus anatinus* (*Oa*), *Monodelphis domestica* (*Md*), *Dasypus novemcinctus* (*Dn*), *Mus musculus* (*Mm*), *Homo sapiens* (*Hs*). Shading represents amino acid groups conserved across the family: *dark green*, hydrophobic and aromatic; *light green*, polar; *blue*, acidic; all glycine and proline residues are highlighted in *yellow*. **(B)** Crystal structure 7LT0 of yeast ScNSE6 and AlphaFold model of moss PpNSE6. The central helical motifs are in dark colours, and the conserved basic contact residues are shown as sticks. **(C** and **D)** Expression of the Gal4AD-PpNSE6 constructs (used in Figs. 1B and C) was verified using the anti-Gal4AD antibody.

**Supplementary Table ST1**

**Supplementary Table ST2**

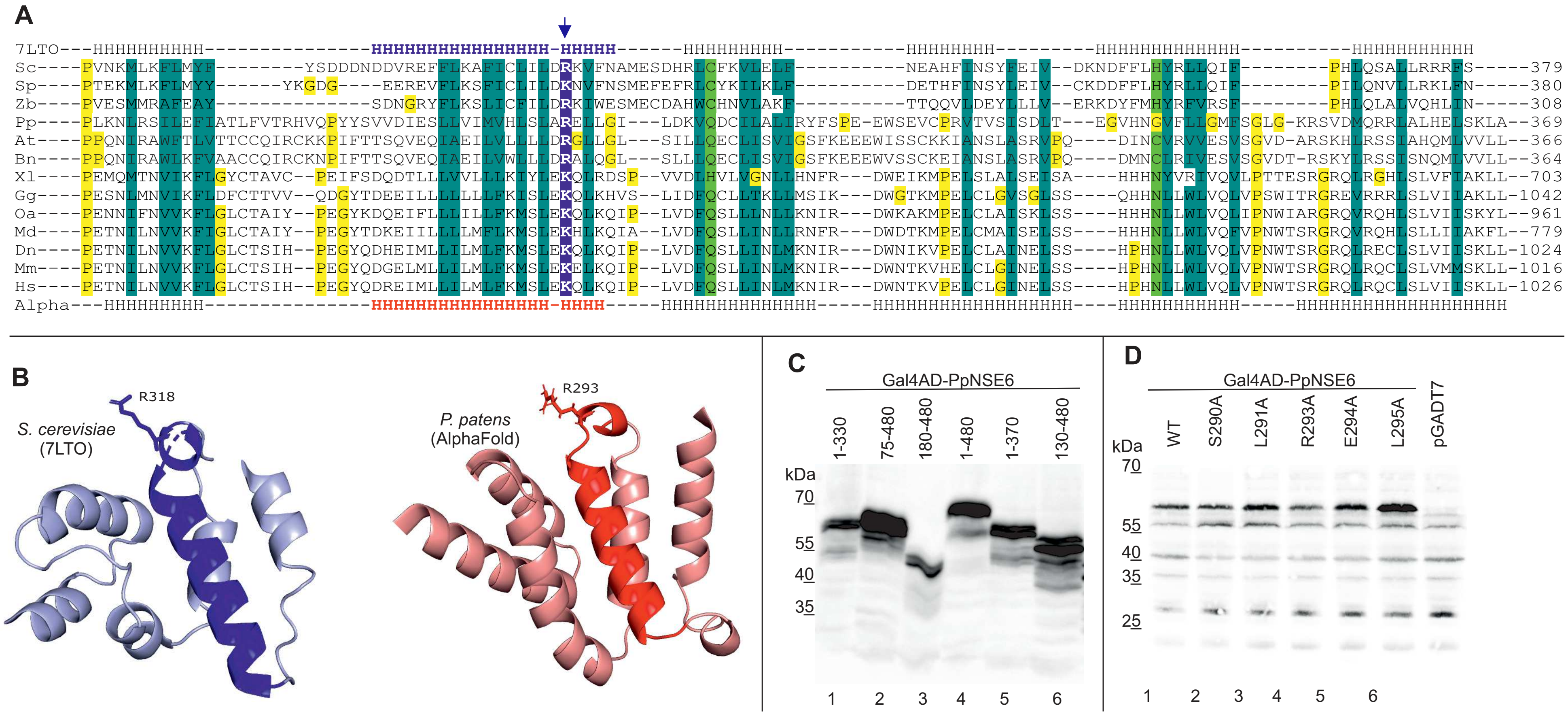

A
7LTO---HHHHHHHHHH----------------HHHHHHHHHHHHHHHHH-HHHHH------HHHHHHHHH----------HHHHHHHH---------HHHHHHHHHHHHH----------HHHHHHHHHHH
Sc----PVNKMLKFLMYF---------YSDDDNDDVREFFLKAFICLILDRKVFNAMESDHRLCFKVLELF-----------NEAHFINSYFEIV--DKNDFFLHYRLLQIF---------PHLQSALLRRRFS-----379
Sp----PTEKMLKFLMYY------YKGDG----EEREVFLKSFICLILDKNVFNSMEFEFRLCYKILKLF-----------DETHFINSYLEIV--CKDDFFLHYRLLQIF---------PILQNVLLRKLFN-----380
Zb----PVESMMRAFEAY----------------SDNGRYFLKSLICFILDRKIWESMECDAHWCHNVLAKF-----------TTQQVLDEYLLLV--ERKDYFMHYRFVRSF---------PHLQLALVQHLIN-----308
Pp----PLKNLRSILEFIATLFVTRHVQPYYSVVDIESLLVIMVHLSLARELLGI---LDKVQDCILALIRYFSPE-EWSEVCPRVTVSISDLT---EGVHNGVFLLGMFSGLG-KRSVDMQRRLALHELSKLA--369
At----PPQNIRAWFTLVTTCCQIRCKKPIFTTSQVEQIAEILVLLLLDRGLLGL---SILLQECLISVIGSFKEEEWISSCKKIANSLASRVPQ----DINCVRVVESVSGVD-ARSKHLRSSIAHQMLVVLL--366
Bn----PPQNIRAWLKFVAACCQIRCKNPIFTTSQVEQIAEILVWLLLDRALQGL---SLLLQECLISVIGSFKEEEWVSSCKEIANSLASRVPQ----DMNCLRIVESVSGVDT-RSKYLRSSISNQMLVVLL--364
Xl----PEMQMTNVIKFLGYCTAVC--PEIFSDQDTLLLVVLLLKIYLEKQLRDSP--VVDLHVLVGNLLHNFR---DWEIKMPELSLALSEISA----HHHNYVRIVQVLPTTESRGRQLRGHLSLVFIAKLL--703
Gg----PESNLMNVIKFLDFCTTVV--QDGYTDEEILLLLLLLFKISLEKQLKHVS--LIDFQCLLTKLLMSIK---DWGTKMPELCLGVSGLSS----QHHNLLWLVQLVPSWITRGREVRRRLSLVIIAKLL-1042
Oa----PENNIFNVVKFLGLCTAIY--PEGYKDQEIFLLLILLFKMSLEKQLKQIP--LVDFQSLLLNLLKNIR---DWKAKMPELCLAISKLSS----HHHNLLWLVQLIPNWIARGRQVRQHLSLVIISKYL--961
Md----PETNILNVVKFIGLCTAIY--PEGYTDKEIILLLMFLKMSLEKHLKQIA--LVDFQSLLINLLRNFR---DWDTKMPELCMAISELSS----HHHNLLWLVQFVPNWTSRGRQVRQHLSLLIIAKFL--779
Dn----PETNILNVVKFLGLCTSIH--PEGYQDHEIMLLILMLFKMSLEKQLKQIP--LVDFQSLLINLMKNIR---DWNIKVPELCLAINELSS----HPHNLLWLVQLVPNWTSRGRQLRECLSLVIISKLL-1024
Mm----PETNILNVVKFLGLCTSIH--PEGYQDGELMLLILMLFKMSLEKELKQIP--LVDFQSLLINLMKNIR---DWNTKVHELCLGINELSS----HPHNLLWLVQLVPNWTSRGRQLRQCLSLVMMSKLL-1016
Hs----PETNILNVVKFLGLCTSIH--PEGYQDREIMLLILMLFKMSLEKQLKQIP--LVDFQSLLINLMKNIR---DWNTKVPELCLGINELSS----HPHNLLWLVQLVPNWTSRGRQLRQCLSLVIISKLL-1026
Alpha---HHHHHHHHH----------------HHHHHHHHHHHHHHHHH-HHHH-----HHHHHHHHHHHHHH-----HHHHHHHHHHHHHHHHHHH---HHHHHHHHHHH-----HHHHHHHHHHHHHHHHHHHH
B
R318
S. cerevisiae
(7LTO)
R293
P. patens
(AlphaFold)
C
Gal4AD-PpNSE6
1-330
75-480
180-480
1-480
1-370
130-480
kDa
70
55
40
35
1
2
3
4
5
6
D
Gal4AD-PpNSE6
WT
S290A
L291A
R293A
E294A
L295A
pGADT7
kDa
70
55
40
35
25
1
2
3
4
5
6

**Supplementary Table ST1 – List of cloning primers**

| | | | |
|---|---|---|---|
| PpNSE6 (aa 1 - 480) to pGADT7 | EB204 | fw | 5′ gacgtaccagattacgctcatatgATGGAGAGCCTTACAACA 3′ |
| | EB207 | rev | 5′ tctgcagctcgagctcgatggatcTCACTGGAAGTTCTCGTCAG 3′ |
| PpNSE6 (aa 75 - 480) to pGADT7 | EB206 | fw | 5′ gacgtaccagattacgctcatatgGATGAGGAGGGTGAGG 3′ |
| | EB207 | rev | 5′ tctgcagctcgagctcgatggatcTCACTGGAAGTTCTCGTCAG 3′ |
| PpNSE6 (aa 180 - 480) to pGADT7 | EB208 | fw | 5′ gacgtaccagattacgctcatatgATCGACGAATCTTTGGAAAG 3′ |
| | EB207 | rev | 5′ tctgcagctcgagctcgatggatcTCACTGGAAGTTCTCGTCAG 3′ |
| PpNSE6 (aa 1 - 330) to pGADT7 | EB204 | fw | 5′ gacgtaccagattacgctcatatgATGGAGAGCCTTACAACA 3′ |
| | EB205 | rev | 5′ tctgcagctcgagctcgatggatccTCATGAGATACTCACAGTCACG 3′ |
| PpNSE6 (aa 1 - 370) to pGBKT7 | MP480 | fw | 5′ gaggaggacctgcatatggccATGGAGAGCCTTACAACAG 3′ |
| | MP481 | rev | 5′ ggccgctgcaggtcgacggatccTTATCGAGCGAGCTTCGATAG 3′ |
| PpNSE6 (aa 75 - 370) to pGBKT7 | EB239 | fw | 5′ atctcagaggaggacctgcatatgGATGAGGAGGGTGAGG 3′ |
| | MP481 | rev | 5′ ggccgctgcaggtcgacggatccTTATCGAGCGAGCTTCGATAG 3′ |
| PpNSE6 (aa 130 - 480) to pGBKT7 | MP482 | fw | 5′ gaggaggacctgcatatggccatggagCACCGGGCGTTGCAAGCA 3′ |
| | MP483 | rev | 5′ ggccgctgcaggtcgacggatccTCACTGGAAGTTCTCGTCAGCA 3′ |
| PpNSE5 (aa 1 - 526) to pGBKT7 | MP441 | fw | 5′ atctcagaggaggacctgcatATGCCTCGCAAAAAGCAG 3′ |
| | MP442 | rev | 5′ ggccgctgcaggtcgacggatccTCAGGAAACTTGCAGGGTA 3′ |
| PpNSE5 (aa 109 - 526) to pGBKT7 | EB278 | fw | 5′ atctcagaggaggacctgcatatgATGTCTGTTGTACCAGATGTTG 3′ |
| | EB258 | rev | 5′ gcggccgctgcaggtcgacggatccTCAGGAAACTTGCAGGGTATG 3′ |
| PpSMC5 (aa 201 - 890) to pGBKT7 | 1280 | fw | 5′ gaggaggacctgcatatggccATGGTGACTCTTATTAAGAAAAATG 3′ |
| | 1281 | rev | 5′ atccccgggaattcggcctcTCACTTGGCCTTGACGGAG 3′ |
| PpSMC5 (aa 280 - 790) to pGBKT7 | MP414 | fw | 5′ atctcagaggaggacctgcatATGAAGCGTCTGCTTAATGAAG 3′ |
| | MP415 | rev | 5′ gcggccgctgcaggtcgacggatccTTATTTACTGTCGTCATATTCCCTC 3′ |
| PpSMC5 (aa 360 - 710) to pGBKT7 | MP416 | fw | 5′ atctcagaggaggacctgcatATGATTGTTGCAGCTACCAGAGA 3′ |
| | MP417 | rev | 5′gcggccgctgcaggtcgacggatccTTACTCCGCGTGAATGGACTCC 3′ |
| PpSMC6 (aa 226 - 955) to pGBKT7 | EB259 | fw | 5′ atctcagaggaggacctgcatatgCAACAAGTATCGGACTTGC 3′ |
| | EB260 | rev | 5′ caggtcgacggatccccgggaattcTCAGTTACGTTCAAATTTGCTACAAC 3′ |
| PpSMC6 (aa 290 - 870) to pGBKT7 | MP410 | fw | 5′ atctcagaggaggacctgcatATGAAGTGGGTTCAAATCACC 3′ |
| | MP411 | rev | 5′ caggtcgacggatccccgggaattcTTAGCAAACTTGCAGAGCCTT 3′ |
| PpSMC6 (aa 350 - 810) to pGBKT7 | MP412 | fw | 5′ atctcagaggaggacctgcatATGCAGCTGAGAAACTCTCAGAG 3′ |
| | MP413 | rev | 5′ caggtcgacggatccccgggaattcTTACCGAGAAGCTTCGATGTCC 3′ |
| PpSMC6 (aa 1 - 510) to pGBKT7 | EB241 | fw | 5′ atctcagaggaggacctgcatatgATGCGGAATAACACTCGG 3′ |
| | EB261 | rev | 5′ gaattcggcctccatggccaCCAATGGAGAAATCGCGTTCG 3′ |
| PpSMC6 (aa 655 - 1095) to pGBKT7 | EB244 | fw | 5′ gcggccgctgcaggtcgacggatccTTAAGGACGAGGGGCC 3′ |
| | EB262 | rev | 5′ atggaggccgaattcccgggGGGAGTGAGACTGTATTGC 3′ |
| Ppnse6/S290A mutation | EB324 | fw | 5′ GATCATGGTGCACCTTgcCCTTGCTCGGG 3′ |
| | EB325 | rev | 5′ CCCGAGCAAGGgcAAGGTGCACCATGATC 3′ |
| Ppnse6/L291A mutation | EB326 | fw | 5′ CATGGTGCACCTTAGCgcTGCTCGGGAATTGCTTG 3′ |
| | EB327 | rev | 5′ CAAGCAATTCCCGAGCAgcGCTAAGGTGCACCATG 3′ |
| Ppnse6/R293A mutation | EB275 | fw | 5′ GGTGCACCTTAGCCTTGCTgcGGAATTGCTTGGAATATTG 3′ |
| | EB276 | rev | 5′ CAATATTCCAAGCAATTCCgcAGCAAGGCTAAGGTGCACC 3′ |
| Ppnse6/E294A mutation | EB328 | fw | 5′ CTTAGCCTTGCTCGGGcATTGCTTGGAATATTG 3′ |
| | EB329 | rev | 5′ CAATATTCCAAGCAATgCCCGAGCAAGGCTAAG 3′ |
| Ppnse6/L295A mutation | EB330 | fw | 5′ CCTTAGCCTTGCTCGGGAAgcGCTTGGAATATTGGAC 3′ |
| | EB331 | rev | 5′ GTCCAATATTCCAAGCgcTTCCCGAGCAAGGCTAAGG 3′ |

### Supplementary Table ST2

| | | | |
|---|---|---|---|
| sgRNA PpNSE6, CRISPRi | pKA1355 | fw | 5′ ATTGAGCCTTACAACAGAGGATG 3′ |
| | pKA1356 | rev | 5′ AAACCATCCTCTGTTGTAAGGCT 3′ |
| sgRNA PpNSE6, knock-in | pKA1402 | fw | 5′ ATTGCGACGAATCAACTTTACGT 3′ |
| | pKA1403 | rev | 5′ AAACACGTAAAGTTGATTCGTCG 3′ |
| DNA donor template | pKA1400 | fw | 5′ GTAGATGGAGAcgCTTACAACAGAGGATtAGAGGACTGGC 3′ |
| | pKA1401 | rev | 5′ GCCAGTCCTCTaATCCTCTGTTGTAAGcgTCTCCATCTAC 3′ |
| PpNSE6, control | pKA1451 | fw | 5′ GCGATGTTGTGCGTAGAACC 3′ |
| | pKA1452 | rev | 5′ CCTTGGCCTTCAATTTCTGG 3′ |
| PpNSE6, transcript | pKA1290 | fw | 5′ CGCAGTTAGAACCTCCATCC 3′ |
| | pKA1291 | rev | 5′ GGCTAAACCCTCCAAAGAAG 3′ |
| qPCR CAP-50 | pKA1001 | fw | 5′ TACAATGCGGCTACCGAC 3′ |
| | pKA1002 | rev | 5′ AGGCCGGATGCAGTAAAC 3′ |
| qPCR 18S rDNA | Pp18S A | fw | 5′ CCTCTAAGAAGTTGGCCGCA 3′ |
| | Pp18S B | rev | 5′ GGCCGTTCTTAGTTGGTGGA 3′ |
| qPCR 5S rDNA | Pp5S F | fw | 5′ TACCAAGGCTACTACACCAGATC 3′ |
| | Pp5S R | rev | 5′ AGGTCACCCATCCCAGTACTA 3′ |
| qPCR ubiquitin | ubqFw | fw | 5′ ACTACCCTGAAGTTGTATAGTTCGG 3′ |
| | ubqRev | rev | 5′ CAAGTCACATTACTTCGCTGTCTA 3′ |
